# Climate warming undermines the benefits of grazing removal for alpine plant population maintenance on the Tibetan Plateau

**DOI:** 10.64898/2026.07.18.739315

**Authors:** Hai-Tao Miao, Yu-Kun Hu, Zhenhua Zhang, Xiaofang Wang, Jin-Sheng He, Shou-Li Li

**Affiliations:** State Key Laboratory of Herbage Improvement and Grassland Agro-Ecosystems, Center for Grassland Microbiome, and College of Pastoral Agriculture Science and Technology, Lanzhou University, Lanzhou, China; State Key Laboratory of Herbage Improvement and Grassland Agro-Ecosystems, College of Pastoral Agriculture Science and Technology, Lanzhou University, Lanzhou, China; Qinghai Haibei National Field Research Station of Alpine Grassland Ecosystem and Key Laboratory of Adaptation and Evolution of Plateau Biota, Northwest Institute of Plateau Biology, Chinese Academy of Sciences, Xining, China; Center for Grassland Microbiome, College of Pastoral Agriculture Science and Technology, Lanzhou University, Lanzhou, China; Institute of Ecology, College of Urban and Environmental Sciences, Peking University, Beijing, China

**Keywords:** alpine plants, demographic compensation, grazing removal, long-term climate warming, stochastic population dynamics

## Abstract

A central question in restoring degraded grasslands is whether grazing removal can sustain viable plant populations under both current and future warming conditions. Addressing this question requires demographic studies integrating vital rates’ responses to grazing removal and climate warming throughout a species’ life cycle. However, studies of this nature are rare. Using stochastic Integral Projection Models parameterized with four years (2020−2023) of demographic data, we find that nine years of grazing removal increases the stochastic population growth rate (logλ_S_) of two coexisting herbaceous plants, *Carex atrofusca* and *Sibirotrisetum sibiricum* at two altitudes (3,700 m and 4,000 m) in an alpine grassland on the Tibetan Plateau. Although individual survival declines following grazing removal, these negative effects are overcompensated by enhanced plant growth, ultimately promoting logλ_S_ in both species. However, the benefits of grazing removal are cancelled under nine years of *in situ* active warming (+2□), where no demographic compensation occurred (*i*.*e*., vital rates change in the opposite directions among populations), and logλ_S_ are even lower than those under grazing. Our findings suggest that while grazing removal is a sustainable management strategy under current climate conditions, it may not remain effective under projected warming, providing valuable information for sustainable population management under global change.

## Introduction

Livestock grazing is a major anthropogenic disturbance to global biodiversity (Watkinson & Ormerod 2001; Gao & Carmel 2020). Overgrazing poses a severe threat to species’ viability worldwide (Hejcman et al. 2013; Bardgett et al. 2021; Sun et al. 2022). To protect species in danger, grazing removal has been extensively implemented as a conservation management practice (Verdoodt et al. 2009; Golodets et al. 2010; Wang et al. 2022; Schrama et al. 2023). However, quantitatively assessing the extent to which grazing removal can shape population viability is challenging, as it requires demographic studies integrating vital rates (*e*.*g*., survival, growth, and reproduction) over an entire life cycle in response to long-term grazing removal. Furthermore, grazing removal may alter multiple vital rates simultaneously, with the magnitude and direction of these effects varying according to the environments inhabited by the populations (Jónsdóttir 1991; Maron et al. 2014; Larios & Hallett 2022). However, we still lack mechanistic insights into impacts of such vital rates’ responses to grazing removal on the long-term population viability. This knowledge gap hinders our ability to design effective grazing management to protect populations of conservation concern.

Whether species can maintain viable populations under grazing removal may depend on long-term climate warming (Zhong et al. 2019; Wang et al. 2022). It may take multiple years for populations, especially populations of perennials, to respond to elevated temperature (Tenhumberg et al. 2018; Evers et al. 2021; Miao et al. 2025). Existing findings about the impacts of climate warming on the viability of ungrazed population are inconsistent, with negative and positive effects being documented at the demography in a few demographic studies (Williams et al. 2007; Evju et al. 2010, 2011; Sletvold et al. 2013). For example, Evju et al. (2010) reported that elevated temperatures in July decreased both survival and growth of the small herb *Viola biflora* in an alpine habitat, thereby accelerating its population decline following cessation of grazing. Conversely, Sletvold et al. (2013) found that elevated summer temperatures enhanced survival and growth of the rare orchid *Dactylorhiza lapponica* in the boreal zone, thereby promoting its population growth under grazing removal. Additionally, Williams et al. (2007) demonstrated that four years of warming by 2°C reduced the population growth of four coexisting perennial plant species under grazing removal in southeastern Australia. One reason for such inconsistent findings may be that most results are based on short-term warming scenarios (*e*.*g*., lasting a month, a season, or less than five years), which only capture the transient dynamics of ungrazed populations under climate warming. However, plant populations may take multiple years, or even decades, to fully respond to climate warming (Tenhumberg et al. 2018; Evers et al. 2021; Miao et al. 2025). Furthermore, it is unclear to what extent warming effects on population dynamics influence the outcomes of grazing removal. These knowledge gaps hamper our ability to prospectively assess the impact of grazing removal on population dynamics in the face of warming climate.

The effects of grazing removal on population viability under both ambient and warmed conditions are inherently determined by the integrated response of all the demographic rates. Grazing removal may affect demographic rates differently, not only in magnitude but also in direction (Jónsdóttir 1991; Maron et al. 2014; Larios & Hallett 2022). When declines in some demographic rates are offset by increases in others, the overall population-level effects of grazing removal could be alleviated or cancelled out, a phenomenon known as “demographic compensation” (Villellas et al. 2015; Sheth & Angert 2018). For instance, Evju et al. (2010) reported that the alpine plant *Viola biflora* compensated for reduced growth through increases in fecundity and stasis, thereby alleviating population decline under grazing removal. In contrast, populations lacking demographic compensation are more prone to decline under grazing removal (Li et al. 2013; Larios & Hallett 2022). Furthermore, the extent to which population persistence is affected by grazing removal depends not only on the magnitude of changes in vital rates but also on the sensitivity of population growth to these changes (de Kroon et al. 2000; Villellas et al. 2015). Consequently, comprehensive modeling that accounts for the cumulative effects of all demographic rates is essential for accurately assessing population viability. Evaluating the presence of demographic compensation under both ambient and warmed conditions is crucial for predicting population responses and informing adaptive grazing management under climate change.

Alpine grasslands are among the most vulnerable ecosystems to grazing activity and climate warming (Körner 2007; Elsen & Tingley 2015; Seddon et al. 2016). The Tibetan Plateau, often referred to as the “Third Pole,” has undergone decades of intensive overgrazing coupled with a rapid warming rate—nearly twice the global average (Wang et al. 2022; Yao et al. 2022). Therefore, understanding the demographic consequences of grazing removal and climate warming on the Tibetan Plateau is crucial for informing the management of such critical susceptible alpine ecosystems. Moreover, such insights can offer early warning signals for the conservation of other ecosystems. While previous studies have found that grazing removal increases aboveground biomass in alpine grasslands on the Tibetan Plateau, such positive effects diminish with altitude (Wang et al. 2022; Xiang et al. 2023). However, it remains unclear how alpine plant population maintenance will respond to grazing removal and climate warming, and whether such effects will be weakened at higher altitudes.

Here, we assess the effectiveness of grazing removal in both ambient and warmed condtions for two coexisting alpine species, *Carex atrofusca* and *Sibirotrisetum sibiricum* on the alpine grassland of the Tibetan Plateau. We tracked the population dynamics of both species in grazed, ungrazed and ungrazed + warmed plots across four years (2020-2023) at different altitudes (3,700 m versus 4,000 m). We utilized these demographic data to parameterize stochastic integral projection models (stochastic IPMs) (Rees & Ellner 2009; Ellner et al. 2016). We conducted a small noise approximation life table response experiment (SNA-LTRE) to decompose differences in stochastic population growth rates into contributions from differences in means, elasticities, variability and correlations in vital rates (Davison et al. 2013; Davison et al. 2019). We used stochastic population models to test the following hypotheses: (1) grazing removal will positively affect vital rates and subsequently increase the long-term population growth rate, because alpine plants are generally favored by herbivory removal. However, the positive effects are expected to diminish towards higher altitude, as the beneficial effects of grazing removal on alpine plants weaken with altitude. (2) Climate warming under grazing removal will negatively affect vital rates and consequently reduce the long-term population growth rate, because alpine plants are generally vulnerable to elevated temperature. Moreover, the negative effects are predicted to exacerbate towards higher altitude, as alpine plants exhibit greater vulnerability at higher elevations. (3) Overall, the negative effects of climate warming under grazing removal on population growth will outweigh the positive effects of grazing removal, because climate change is a dominant driver of grassland change on the Tibetan Plateau (Zhong et al. 2019; Wang et al. 2022).

## Materials and Methods

### Study area and experimental design

The study was conducted at the Qinghai Haibei National Field Research Station of Alpine Grassland Ecosystem, located in the northeastern of the Tibetan Plateau, Qinghai Province, China (37°36′ N, 101°19′ E; Figure 1a). The region experiences a typical alpine climate characterized by short, cool summers and long, cold winters. The mean annual temperature is −1.1 □, with the warmest month (June) averaging 10.4 □ and the coldest month (January) averaging −14.6 □. The mean annual precipitation is 481 mm, over 80% of which falls during the growing season (May to September), primarily as rainfall. The study area is covered by alpine meadow, with the vegetation dominated by perennial herbaceous species (Figure 1a). Livestock husbandry is the primary land use in the region due to the extensive rangeland area. The study sites have experienced a long grazing history by sheep and cattle. However, improper grazing management, particularly overgrazing resulting from rangeland privatization, is believed to cause severe rangeland degradation or even desertification (Wang et al. 2022).

**Figure 1.**
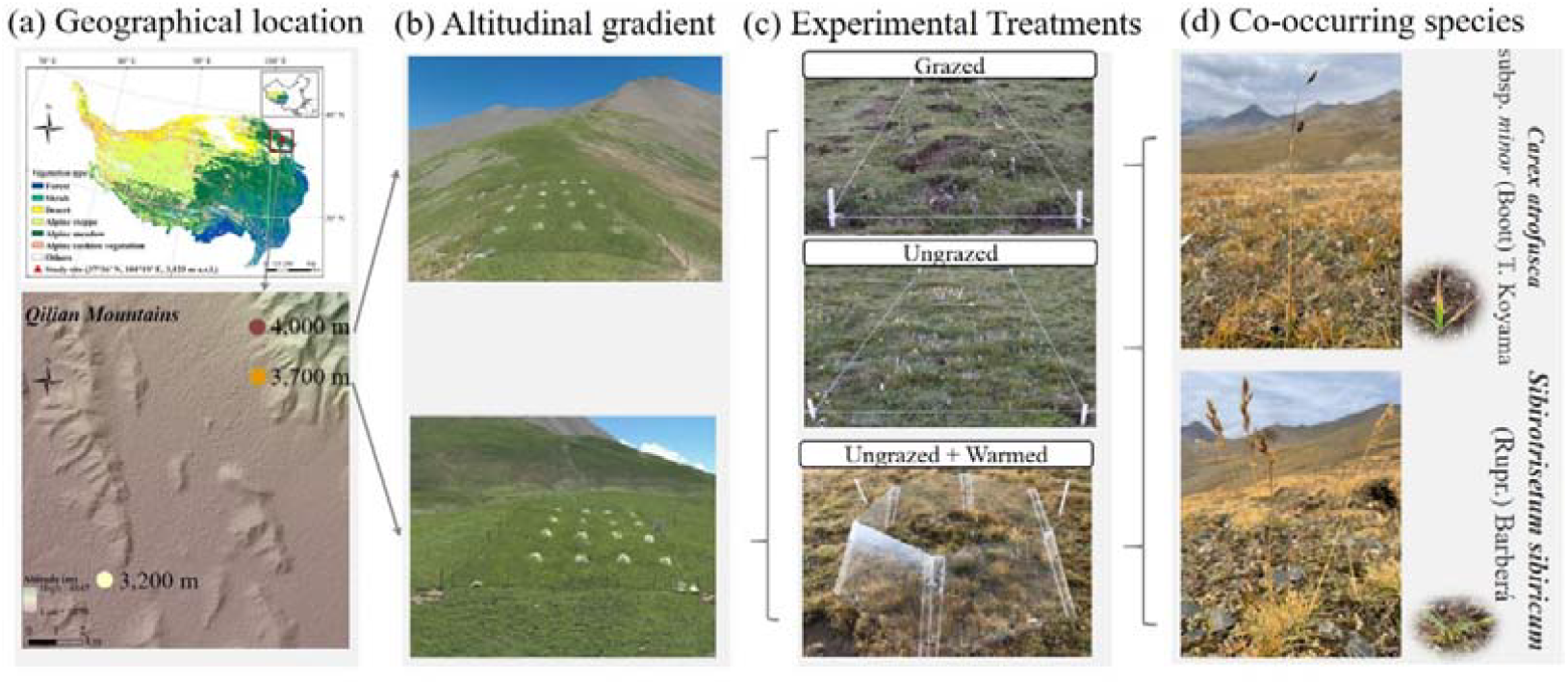
Two co-occurring alpine herbaceous species, *Carex atrofusca* subsp. minor (Boott) T. Koyam and *Sibirotrisetum sibiricum* (Rupr.) Barberá under a long-term grazing removal and warming experiment along an altitudinal gradient (3,700 m and 4,000 m) on the Tibetan Plateau. (a) The geographic location and altitude of the study site on the Tibetan Plateau. (b) The altitudinal gradient includes 3,200 m, 3,700 m and 4,000 m. As the individual number of our study species at 3,200 m was insufficient for demographic study, we focused solely on sites at 3,700 m and 4,000 m. (c) The experimental treatments at each altitude include grazed, ungrazed and ungrazed + warmed. (d) The study species, *C. atrofusca* and *S. sibiricum*, at the study site.

To assess the effects of grazing removal and climate warming on alpine plants, a manipulative field experiment was established along an altitudinal gradient (3,200 m, 3,700 m and 4,000 m) during 2014-2015 (Figure 1b; Table S1). At each altitude, three treatments were implemented: grazed, ungrazed and ungrazed + warmed (Figure 1c), with 3-6 replicate plots per treatment (Table S2). Grazed plots were situated in ambient areas subjected to ongoing livestock grazing by cattle and sheep. Adjacent to these plots, ungrazed plots were fenced with 1.8 m high wire mesh to exclude livestock. To simulate climate warming under grazing removal, transparent hexagonal open-top chambers (OTCs) were installed in the ungrazed plots. Each OTC had an internal diameter of 1.5□m and a height of approximately 0.65□m. This design is commonly used in remote alpine ecosystems to passively elevate temperature (Elmendorf et al. 2012; Prager et al. 2022). The OTCs increased the mean air temperature by ~2□ and soil temperature in the top 5 cm layer by ~1□ (Prager et al., 2022), consistent with the predicted warming of 2□ by 2050 (IPCC 2023). Notably, the warming treatment was only applied within the ungrazed plots to avoid confounding effects of livestock trampling on the OTC structure.

### Study species and demographic data

To compare the effects of grazing removal and climate warming on the population performance of alpine plant species inhabiting the same habitats, we selected two co-occurring alpine species: *Carex atrofusca* subsp. minor (Boott) T. Koyam and *Sibirotrisetum sibiricum* (Rupr.) Barberá. Both species are common species in our study area but differed in taxonomic family and reproductive traits. The plant of *C. atrofusca* is a perennial sedge with flattened basal leaf blades at the base and 2~5 oval spikes borne terminally on a single erect culm at the reproductive stage (Figure 1d). The plant of *S. sibiricum* is a perennial herb with flattened leaf blades at the base and 1-30 loose panicles borne terminally on erect culms at the reproductive stage (Figure 1d). Despite their morphological differences, both species share a similar life history: they sprout new leaves annually, regrow in late April, develop flowering stems from the leaf base in May, flower in late June, set fruit from July to August, and their aboveground parts wither in winter. Recruitment in both species primarily occurs through seed germination in the following spring. The plant of *C. atrofusca* propagates vegetatively via horizontally spreading rhizomes that produce new ramets, while *S. sibiricum* exhibits strong tillering, with tillers capable of developing into independent individuals. The plant of *C. atrofusca* has a broad distribution across the Arctic and the alpine regions of Central Asia, Europe and North America (Schonswetter et al. 2006), thriving at elevations from 2,200 to 4,600 m. The plant of *S. sibiricum* is widespread across temperate regions of Europe and Asia, occurring between 750 and 4,200 m in elevation. Both species are ecologically important as forage plants in alpine grasslands.

To examine the effects of grazing removal and climate warming on demographic processes, we conducted annual censuses of both focal species in August from 2020 to 2023. In the first census, we measured plant height of both species, which served as a proxy for individual-level state variable (Struckman et al. 2019; Miao et al. 2025). We also recorded the reproductive status of each individual for both species in each plot and counted the number of spikes or panicles produced by each reproductive individual. Upon the first measurement, we tagged each individual with a stainless steel label and recorded its Cartesian coordinates within the plots to allow tracing through time. In the subsequent annual census from 2021 to 2023, we checked the survival status of all tagged individuals, remeasured plant height, and recorded the aforementioned reproductive characteristics on surviving ones. We also located and measured new recruits within the plots. Since reproductive individuals of both species typically produce large quantities of seeds, and it is impossible to distinguish whether new recruits originate from seeds or from vegetative propagation (ramets or tillers), we assumed that reproduction was generally size-dependent. Specifically, the number of seeds and vegetative offspring produced by an individual was assumed to be linearly related to its size (Verburg et al. 1996). Accordingly, the number of new recruits produced per individual was estimated by multiplying its size (*x*) by the ratio *n*/∑*x*, where *n* is the total number of new recruits in the current year and ∑*x* is the sum of the sizes of all individuals in the previous year within each plot (Li et al. 2013; Li et al. 2015). In total, we tracked 1,904 individuals of *C. atrofusca* and 1,395 individuals of *S. sibiricum* across all plots located at 3,700 m and 4,000 m (Table S2). Because the number of individuals at 3,200 m was insufficient for demographic analysis, we focused solely on the sites at 3,700 m and 4,000 m.

### Environmental variation

To evaluate how environmental variation affect population dynamics under different grazing intensities, each census period (*i*.*e*., environmental condition) were classified into one of three environmental states: dry, normal, and wet years. This classification was based on total annual precipitation, a key proxy for plant population dynamics (van de Pol et al. 2016; Wang et al. 2020). Specifically, dry years had total precipitation lower than 1 standard deviation, while wet years had total precipitation greater than 1 standard deviation (Xu et al. 2020). To accurately capture the demographic responses to environmental variation, total annual precipitation was calculated by summing the precipitation from October at year *t* to September at year *t + 1* (van de Pol et al. 2016; Wang et al. 2020). Based on the long-term precipitation records spanning 1980-2023, normal years were observed from 2020 to 2021, followed by a wet year in 2021-2022 and a dry year in 2022-2023 (Appendix S1: Figure S1). To incorporate temporal autocorrelation among environmental conditions into stochastic simulations of population dynamics, we defined a discrete Markov chain consisting of three environmental states. Transition probabilities between states were estimated from long-term precipitation records (Appendix S1: Figure S2).

### Demographic rate estimates

To assess the effects of grazing removal and climate warming on demographic processes along an altitudinal gradient, we employed regression models to relate each demographic rate to the log-transformed plant height and treatments at each altitude. Demographic data were collected under each treatment at each altitude over four annual censuses (three transitions), resulting in a total of 18 populations for *C. atrofusca* and 15 for *S. sibiricum*. These data were then pooled and used to fit regression models for five vital rates that collectively determine population dynamics: (a) survival, (b) size changes (*i*.*e*., growth and shrinkage), (c) probability of reproduction, (d) fecundity, and (e) recruit size. Probability of survival and reproduction were modeled from the Binomial family with the logit link function. Size changes and recruit size were modeled from the Gaussian family. Fecundity was modeled from the beta family with the logit link function (Cribari-Neto & Zeileis 2010).

To identify the best predictors of each demographic rate, we constructed multiple regression models incorporating fixed effects of the log-transformed plant height, treatments, and their interaction, as well as random effects of year and plot (Table S3; Ellner et al. 2016; Tredennick et al. 2018). Models were fitted using the R packages *lme4* and *glmmTMB* (Bates et al. 2015; Brooks et al. 2017). We compared candidate models using Akaike’s Information Criterion corrected for small sample size (AICc) to select the best-supported model for each demographic rate (Table S4; Burnham & Anderson 2004). To evaluate the fixed effects in the best-supported model of each demographic rate, Wald chi-squared tests were conducted using the R packages ‘*car*’ (Fox & Weisberg 2019), and significance was determined using type-II tests (Table S5). To evaluate the random year effects in the best-supported model, Likelihood ratio tests (LRT) were performed by comparing models with and without the year term (Yang 1998), with significance determined using Chi-squared test (Table S5).

### Population dynamics model

To quantify the effects of grazing removal and climate warming on long-term population dynamics of alpine species, we constructed stochastic Integral Population Model (IPMs) of both study species (Rees & Ellner 2009). IPMs use information on how an individual’s state influences vital rates to project population change in discrete time (Easterling et al. 2000). Each IPM was parameterized with multi-year estimates of vital rates derived from the best-supported vital rate models. In our IPMs, the continuous state variable was the log-transformed plant height log_*e*_(height) and the discrete time step (from *t* to *t*+ 1) corresponded to one year. The time-varying size-structed IPM is

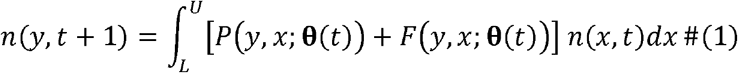

where *L* and *U* are the lower and upper bounds on the range of possible heights log_*e*_(height) for all treatments, and **θ**(*t*) is a vector of year-specific parameters, assumed to vary over time. The variable *n*(*x,t*) is the distribution of log_*e*_(height) *x* at time *t*, and *n*(*y,t* + 1) is the distribution of log_*e*_(height) *y* at time *t*+ 1. The expression *P*(*y,x*; **θ**(*t*)) + *F*(*y,x*; **θ**(*t*)) is called the kernel, *K*(*y,x*; **θ**(*t*)), which is a non-negative surface describing all possible demographic transitions from log_*e*_(height) *x* at time *t* to log_*e*_(height) *y* at time *t*+ 1. The function *P*(*y,x*; **θ**(*t*)) comprises the survival-growth component of a time-varying IPM and can be decomposed into two functions that determine the probability of survival of an individual at log_*e*_(height) *x, S*(*x*; **θ**(*t*)), and the likelihood that the individual will grow from log_*e*_(height) *x* to log_*e*_(height) *y* over a year, *G*(*y,x*; **θ**(*t*)), such that *P*(*y,x*; **θ**(*t*)) = *S*(*x*;**θ**(*t*)) *G*(*y,x*; **θ**(*t*)). The function *F*(*y,x*; **θ**(*t*)) comprises reproductive component of a time-varying IPM and can be decomposed into three functions that determine the probability of reproduction of an individual at log_*e*_(height) *x, F*_0_(*x*; **θ**(*t*)), the number of new recruits produced per individual at log_*e*_(height) *x*, *F*_1_ (*x*; **θ**(*t*)) and the probability distribution of recruit size *F*_2_ (*y*; **θ**(*t*)), such that *F*(*y,x*; **θ**(*t*)) = *F*_*0*_ (*x*; **θ**(*t*)) *F*_1_(*x*; **θ**(*t*)) *F*_2_(*y*; **θ**(*t*)).

To assess the effects of grazing removal and climate warming on the long-term population persistence, we calculated the stochastic population growth rate (logλ_S_) for both species under each treatment and altitude. The logλ_S_ is given by the long-run geometric mean of annual growth rates (Rees & Ellner 2009):

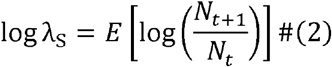

where *N*_*t*_ is the total population size summed across sizes at time *t* and *E* represents the expected value. The logλ_S_ indicates whether a population will increase (logλ_S_ > 0) or decline to extinction (logλ_S_ < 0) for a long time (Tuljapurkar 1990). We first discretized the time-varying IPM kernel into three large transition matrices of size 30 × 30 using the midpoint rule (Ellner & Rees 2006). Using a sequence of Markovian environmental states derived from these matrices (Appendix S1: Figure S2), we then simulated population dynamics for 10,000 years. We then calculated the geometric mean growth rate of the latter *T* = 9,000 years (discarding the first 10% as transient dynamics; Elderd & Miller 2016). To estimate the uncertainty in logλ_S_, we bootstrapped the demographic data 1,000 times for each species to obtain the 95% confidence interval of log,_S_ (Andrello et al. 2020). To examine the effect of grazing removal and climate warming on logλ_S_, we calculated the pairwise difference in logλ_S_ (Δlogλ_S_) between grazing removal and grazing, and climate warming and grazing removal), as well as their 95% confidence intervals. Furthermore, the aforementioned bootstrapped samples were employed in the subsequent population dynamics analysis.

### Small noise approximation life table response experiment

To evaluate how demographic rates contributed to Δlogλ_S_, we performed a small noise approximation life table response experiment (SNA-LTRE; Davison et al. 2013). The Δlogλ_S_ was decomposed into the contributions 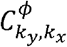, arising from differences in each descriptor *ϕ* (means *μ*; coefficients of variation *c*; temporal correlations *ρ*; elasticities *e*) of stage-specific vital rates *k*_*x*_ (survival *S*_*x*_; shrinkage 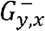; growth 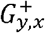; reproduction *F*_0,*x*_; fecundity *F*_1,*x*_; recruit size F_2,*y*_):

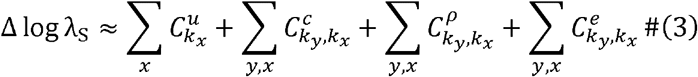

We then derived the contributions 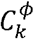 of each vital rate *k* and each descriptor *ϕ* by summing the 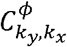 across either stage-specific vital rates or descriptors. To compared relative total effects of each descriptor, we calculated their proportional contributions as 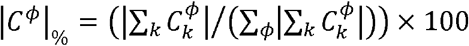 (Davison et al. 2019).

## Results

### The impact of grazing removal and climate warming at different altitudes on demographic processes

To examine the effects of grazing removal and climate warming on the demographic processes of two co-occurring species, *C. atrofusca* and *S. sibiricum*, we used regression models to relate vital rates to log-transformed plant height and treatments at each altitude (3,700 m and 4,000 m). We found that treatments within each altitude and plant size significantly affected all demographic rates of both species, with considerable temporal variation across years (Table S5). The effects of treatments differed among demographic rates, across plant sizes, and between species (Figure 2a, c). Grazing removal consistently decreased the survival of *C. atrofusca* across all plant sizes, whereas it reduced the survival of shorter individuals and increased that of taller individuals in *S. sibiricum* (Figure 2a, c). Under grazing removal, warming reduced the survival of middle individuals in *C. atrofusca* and of shorter individuals in *S. sibiricum* (Figure 2a, c). Grazing removal reduced shrinkage and enhanced growth of middle individuals in both species. Warming under grazing removal had limited effects on changes in plant sizes in *C. atrofusca*, but it increased shrinkage and suppressed growth of taller individuals in *S. sibiricum* (Figure 2a, c). Both grazing removal and warming under exclusion lowered the threshold log-transformed plant height at which individuals reached a 50% probability of reproduction in both species (Figure 2a, c). Grazing removal reduced fecundity but increased recruit size in both species, while warming under grazing removal had minimal effects on these demographic rates (Figure 2a, c).

**Figure 2.**
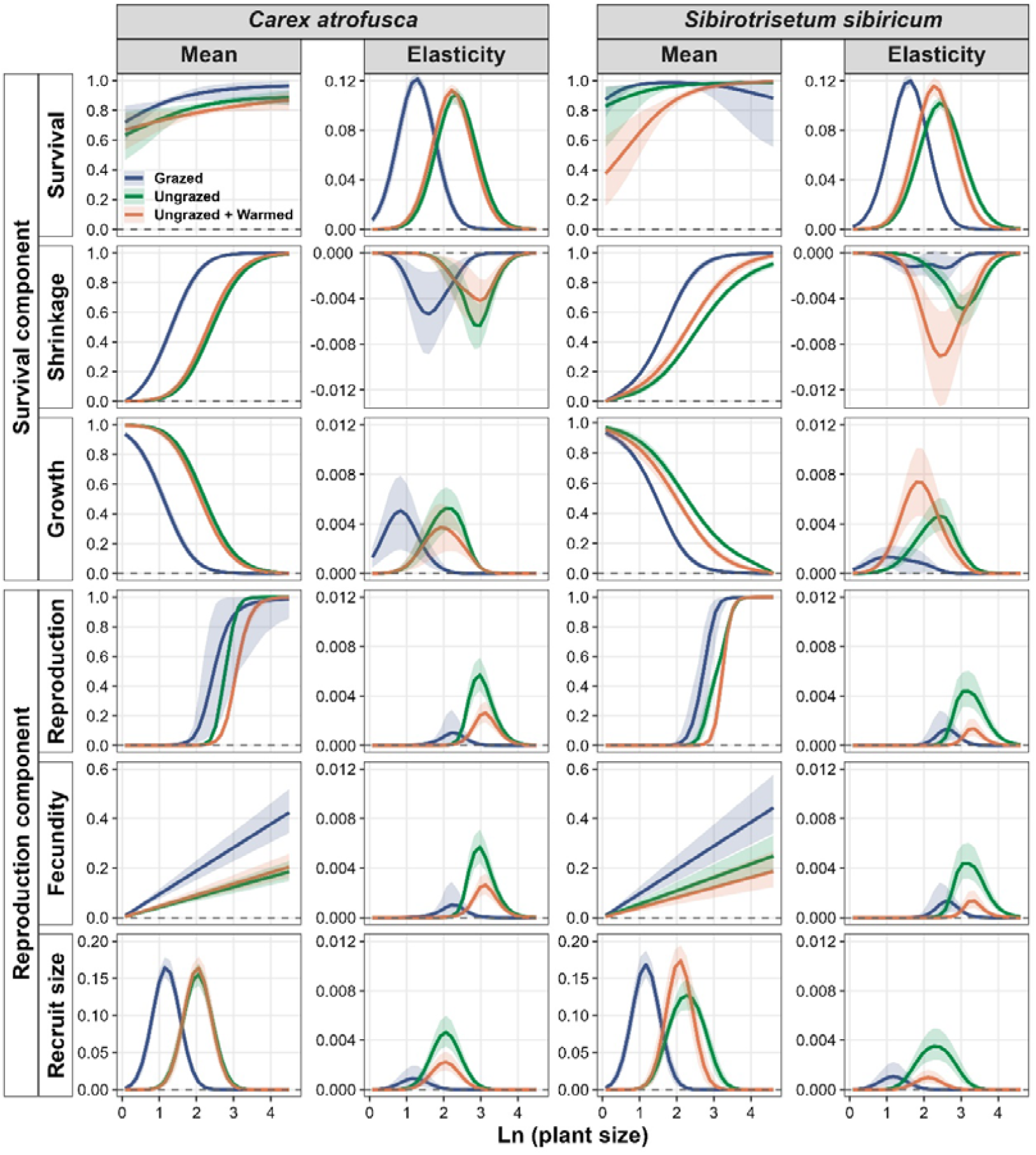
The effects of grazing removal (ungrazed) and warming (ungrazded + Warmed) on mean demographic rates (*i*.*e*., survival, shrinkage, growth, reproduction, fecundity, and recruit size) and their elasticities across the log-transformed plant height for two co-occurring alpine herbaceous species, *Carex atrofusca* and *Sibirotrisetum sibiricum*, on the Tibetan Plateau. Colored lines represent mean values with 95% confidence intervals shown as shaded areas. Both were calculated by randomly sampling statistical over 2,000 bootstrapped demographic datasets.

In addition, increased altitude increased the survival of taller individuals in *C. atrofusca*, but had limited effects on survival in *S. sibiricum* (Figure 3a, c). Increased altitude increased shrinkage and decrease growth of middle individuals in *C. atrofusca*, and of shorter and taller individuals in *S. sibiricum* (Figure 3a, c). Increased altitude raised the threshold log-transformed plant height at which individuals reached a 50% probability of reproduction in both species, while exerting limited effects on fecundity and recruit size (Figure 3a, c). Moreover, altitude modulated the effects of grazing removal and warming on demographic rates (Figure S3a, c, e, g).

**Figure 3.**
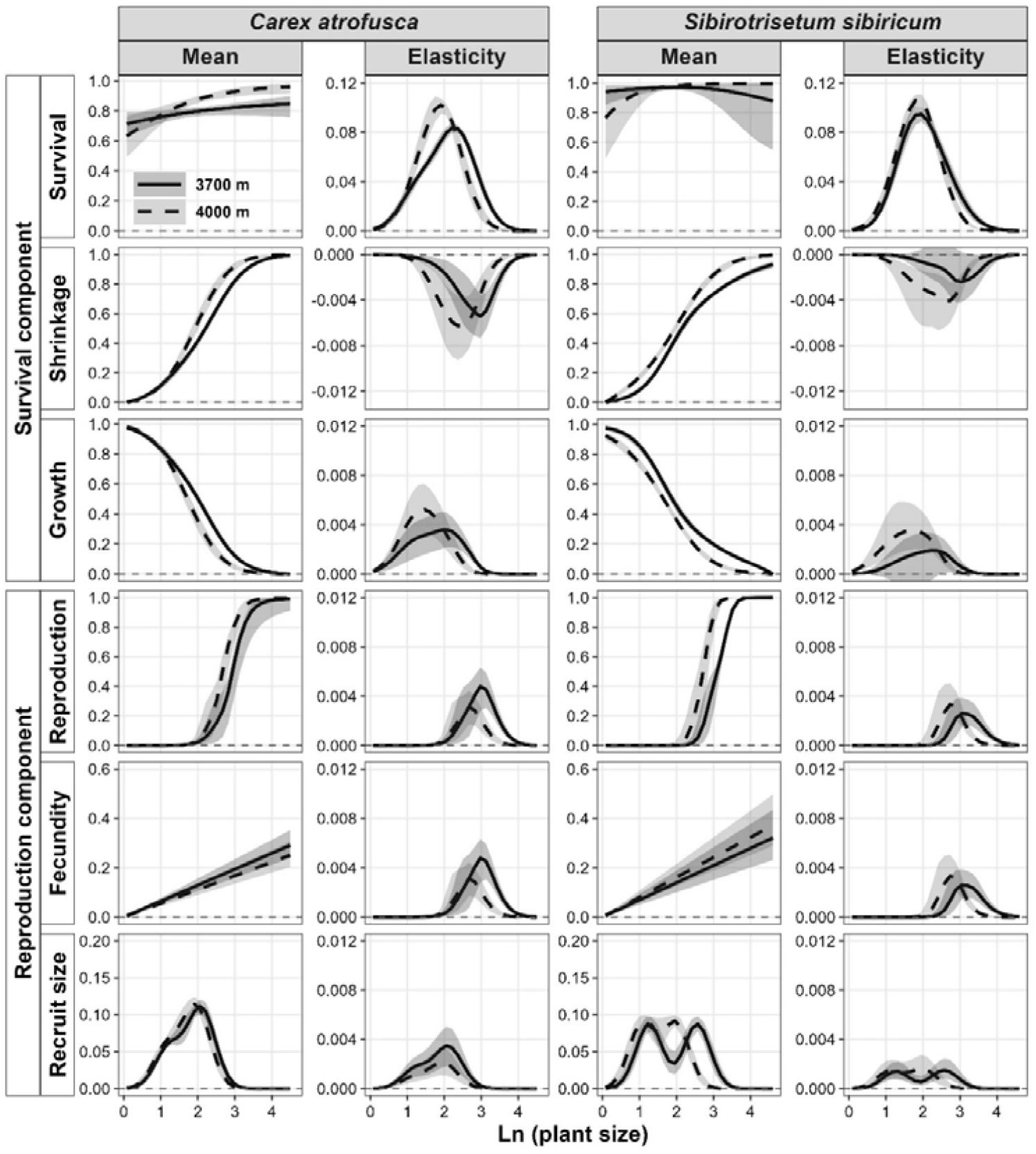
The effects of increased altitude on mean demographic rates (*i*.*e*., survival, shrinkage, growth, reproduction, fecundity, and recruit size) and their elasticities across the log-transformed plant height for two co-occurring alpine herbaceous species, *Carex atrofusca* and *Sibirotrisetum sibiricum*, on the Tibetan Plateau. Lines represent mean values with 95% confidence intervals shown as shaded areas. Both were calculated by randomly sampling statistical over 2,000 bootstrapped demographic datasets.

### The impact of grazing removal and climate warming at different altitudes on the elasticity of demographic processes to population growth

To assess how grazing removal and climate warming affect the relative importance of demographic processes to population maintenance, we calculated the elasticity of each demographic rate to population growth at each altitude (3,700 m and 4,000 m). Across treatments and altitudes, population growth was overwhelmingly most elastic to changes in survival in both species (Figure 2c, d; Figure 3c, d). The elasticities to growth, reproduction, fecundity and recruit size were positive but substantially smaller than that of survival, whereas elasticity to shrinkage was consistently negative, indicating a detrimental effect on population maintenance due to plant size reduction (Figure 2c, d; Figure 3c, d).

In both species, the elasticities increased in shorter individuals, peaked at intermediate plant size, and then declined in taller individuals (Figure 2c, d; Figure 3c, d). Grazing removal shifted the peak elasticity towards taller plants, while warming under grazing removal altered the plant size at which peak elasticity occurred (Figure 2c, d). Increased altitude shifted the peak elasticity towards shorter individuals in both species (Figure 3c, d). Furthermore, altitude modulated the effects of both grazing removal and climate warming on the elasticity of demographic rates (Figure S3b, d, f, h).

### The impact of grazing removal and climate warming under different altitudes on the population growth

To evaluate the effects of grazing removal and climate warming on long-term population persistence, we constructed stochastic integral projection model (stochastic IPM) to estimate the stochastic population growth rate (logλ_S_) of *C. atrofusca* and *S. sibiricum* at two altitudes (3,700 m and 4,000 m). Under grazing conditions, logλ_S_ values differed between species, with *C. atrofusca* exhibiting lower values (logλ_S_ = −0.153, 95% CI [−0.180, −0.126]) than *S. sibiricum* (logλ_S_ = −0.010, 95% CI [−0.020, 0.001]; Figure 4a, d). Grazing removal had a limited positive effect on the population growth of *C. atrofusca* (Δlogλ_S_ = 0.012, 95% CI [−0.025, 0.050]) and significantly increased that of *S. sibiricum* to 0.012 (Δlogλ_S_ = 0.022, 95% CI [0.004, 0.040]; Table S6). In contrast, warming under grazing removal substantially reduced population growth in both species, with logλ_S_ decreasing to −0.210 in *C. atrofusca* (Δlogλ_S_ = −0.069, 95% CI [−0.106, −0.034]) and to −0.104 in *S. sibiricum* (Δlogλ_S_ = −0.116, 95% CI [−0.151, −0.084]; Table 1). Overall, the negative effects of warming on population growth outweighed the positive effects of grazing removal in both species (Table 1).

**Table 1.**
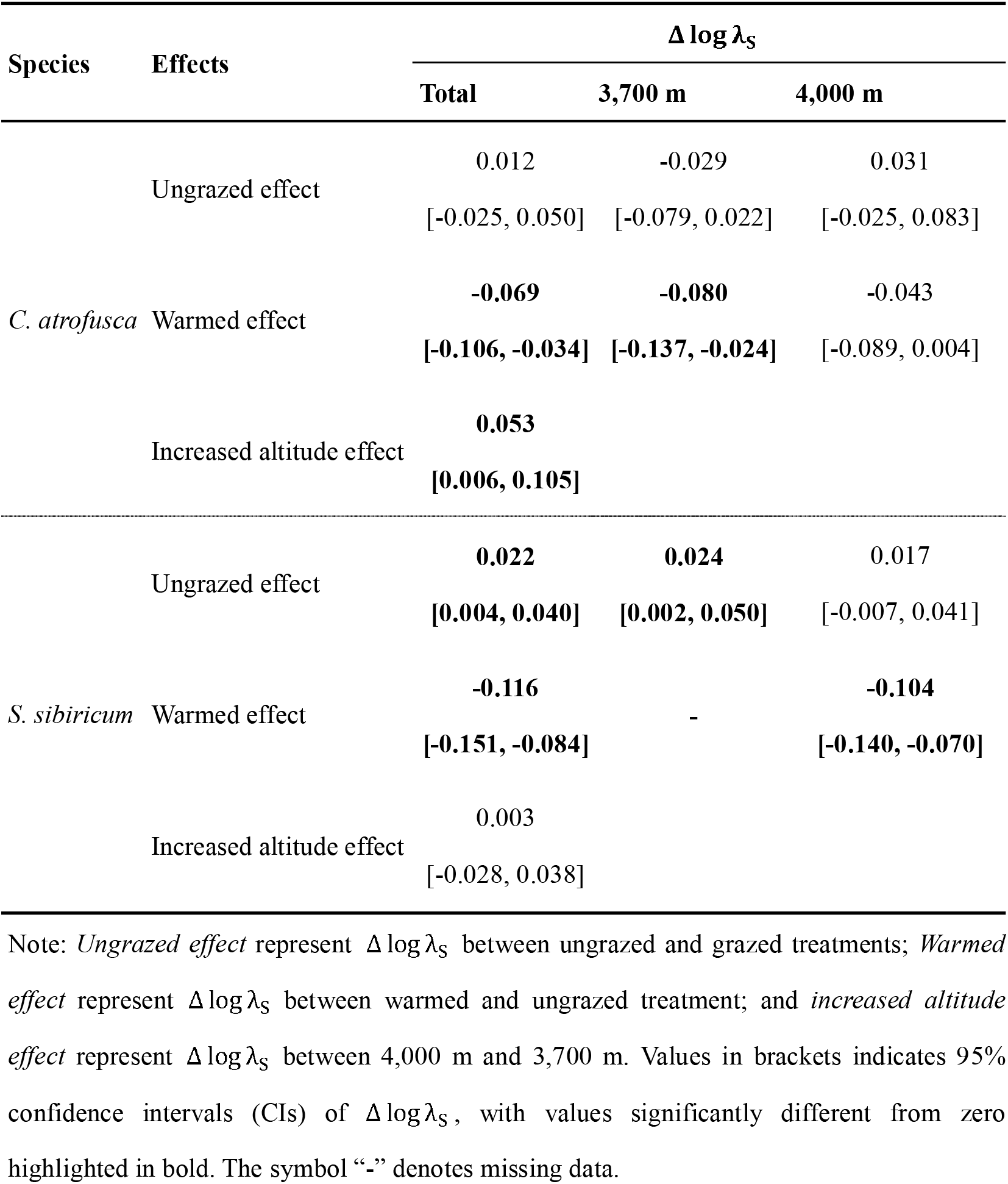
The effects of grazing removal and climate warming on the changes in stochastic population growth rate (Δ log λ_S_) at different altitudes (3,700 m and 4,000 m) for two co-occurring alpine herbaceous species, *Carex atrofusca* and *Sibirotrisetum sibiricum*, on the Tibetan Plateau.

**Figure 4.**
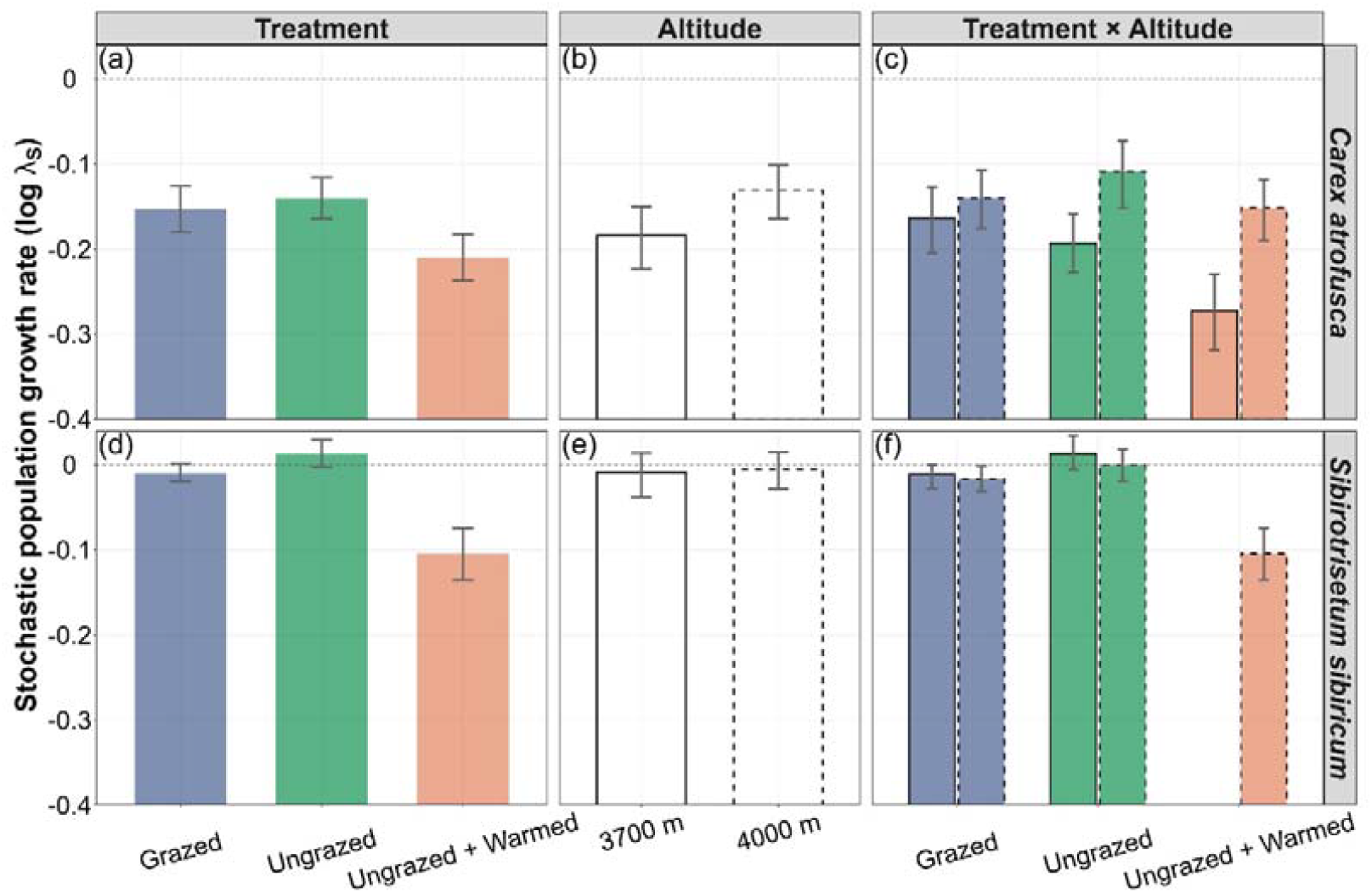
The effects of grazing removal (ungrazed) and warming (ungrazed + warmed) along an altitudinal gradient (3,700 m *vs*. 4,000 m) on the stochastic population growth rate (log λ_S_) for two co-occurring alpine herbaceous species, *Carex atrofusca* (a-c) and *Sibirotrisetum sibiricum* (d-e), on the Tibetan Plateau. Error bars indicate 95% confidence interval of log λ_S_. The horizontal dashed line at zero represents the threshold between population growth (log λ_S_ > 0) and population decline (log λ_S_ < 0).

At 3,700 m, logλ_S_ values differed markedly between species, with *C. atrofusca* showing lower values (logλ_S_ = −0.184, 95% CI [−0.223, −0.150]) than *S. sibiricum* (logλ_S_ = −0.009, 95% CI [−0.038, 0.014]; Figure 4b, e). Increased altitude to 4,000 m promoted the population growth of *C. atrofusca* (Δlogλ_S_ = 0.053, 95% CI [0.006, 0.105]), while it had negligible effect on *S. sibiricum* (Table S6). Importantly, the effects of grazing removal and climate warming on the population growth in both species were modulated by altitude (Figure 4c, f). Grazing removal had a minor negative effect on *C. atrofusca* at 3,700 m but a minor positive effect at 4,000 m (Table 1). In contrast, it increased logλ_S_ of *S. sibiricum* to 0.013 at 3,700 m (Δlogλ_S_ = 0.024, 95% CI [0.002, 0.050]), while exerting a limited positive effect at 4,000 m (Table 1). Warming under grazing removal substantially reduced logλ_S_ of *C. atrofusca* to −0.273 at 3,700 m (Δlogλ_S_ = −0.080, 95% CI [−0.137, −0.024]) but had a small negative effect at 4,000 m. Conversely, warming under grazing removal caused population extinction of *S. sibiricum* at 3700 m and reduced logλ_S_ to −0.104 at 4000 m (Δlogλ_S_ = −0.104, 95% CI [−0.140, −0.070]; Table 1).

### Contributions of demographic rates to changes in population growth under grazing removal and climate warming

To assess how changes in demographic rates contributed to changes in the stochastic population growth rate (Δlogλ_S_) under grazing removal and climate warming, we conducted a small noise approximation life table response experiment (SNA-LTRE) analysis at each altitude (3,700 m and 4,000 m). We found that changes in means of vital rates overwhelmingly contributed to Δlogλ_S_ for both species (97%), whereas stochastic components made relatively small but non-negligible contributions (3%) (Figure S4). The limited increased Δlogλ_S_ of *C. atrofusca* under grazing removal was primarily maintained by compensatory increases in contributions from growth and shrinkage of middle individuals that offset reductions in survival, reproduction, and fecundity (Figure 5a; Figure S5a, c). In contrast, the increased Δlogλ_S_ of *S. sibiricus* under grazing removal was mainly driven by enhanced contributions from growth and shrinkage of middle individuals, despite substantial reductions in survival, reproduction, and fecundity (Figure 5b; Figure S5b). Under warming combined with grazing removal, the decreased Δlogλ_S_ in both species was predominantly attributed to reduced survival of middle individuals, with smaller negative contributions from growth, shrinkage, and reproduction (Figure 5c, d; Figure S5e, h). The increased Δlogλ_S_ of *C. atrofusca* towards higher altitude was largely due to increased survival with a lesser positive contribution from reproduction of middle individuals, despite substantial negative contributions from shrinkage and growth (Figure 5e). Conversely, the unchanged Δlogλ_S_ of *S. sibiricus* towards higher altitude was maintained by compensatory increase in reproduction of middle individuals that offset the negative contribution from shrinkage (Figure 5f).

**Figure 5.**
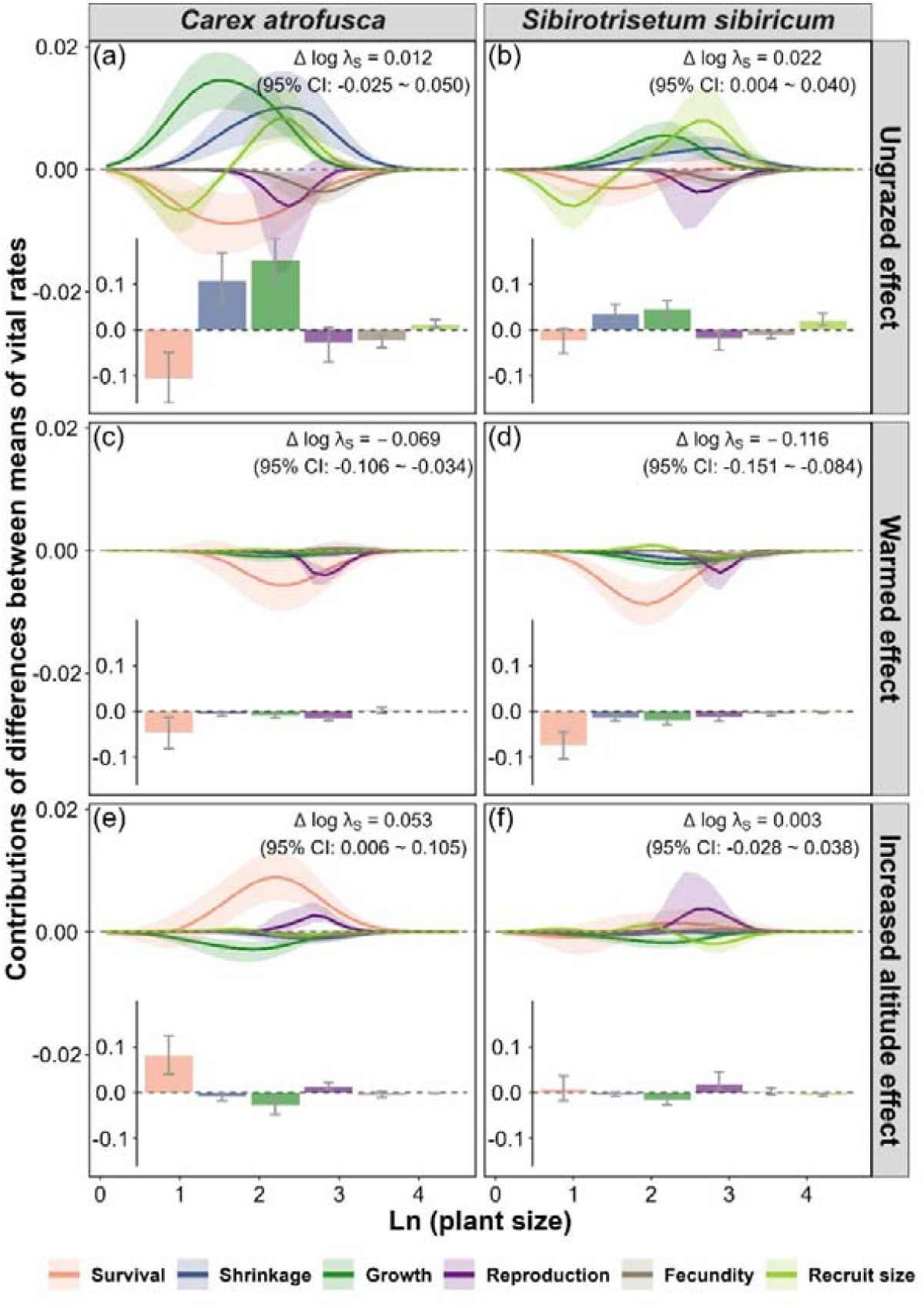
Results of the small noise approximation life table response experiment (SNA-LTRE) to examine the effects of grazing removal, warming, and increased altitude on the contributions of differences between means of vital rates to changes in stochastic population growth rate (Δ log λ_S_) for two co-occurring alpine herbaceous species, *Carex atrofusca* and *Sibirotrisetum sibiricum*, on the Tibetan Plateau. The contribution values of differences between means of each vital rate accumulated through all plant size are given in the bar chart at the bottom of each panel. Shaded areas and error bars indicate 95% confidence intervals of contributions.

## Discussion

### Population dynamics under current grazing in ambient and warmed condtions

Quantitatively assessing the long-term persistence of plant populations under current grazing practices and projected climate warming is crucial for developing optimal grazing management strategies and guiding future biodiversity conservation efforts (Nomoto & Alexander 2021; Larios & Hallett 2022). Comparing the demographic responses of co-occurring species in stochastic environments across multiple altitudes can help generalize findings about the impacts of grazing and climate warming on plant species (Menges 2000; Miao et al. 2025). However, such comparative approaches have been rarely applied in previous demographic studies. Here, we quantified the demographic responses of two coexisting alpine species, *C. atrofusca* and *S. sibiricum*, to grazing removal and to a decade of *in situ* active warming under grazing removal across different altitudes over a four-year period on the Tibetan Plateau. Although current grazing activities lead to contrasting fates for the two species (*C. atrofusca* experiencing population decline and *S. sibiricum* maintaining a stable population), grazing removal facilitated the population growth in both species. The positive effects of grazing removal on population persistence of herbaceous species have been widely documented across diverse ecosystems (Knight 2004; Elderd & Doak 2006; Martorell 2007; Farrington et al. 2009; Bermingham 2010). For example, Elderd & Doak (2006) found that herbivore exclusion alleviated the population decline of the common plant *Minmulus guttatus* in riparian habitats of California; Martorell (2007) reported fencing stabilized the declining population of the threatened herb *Echeveria longissimi* in degraded steep hillside of Mexico; Knight (2004) found that herbivory removal reversed the declining population of the perennial herb *Trillium grandiflorum* to growth within an old-growth beech and maple forest; Farrington et al. (2009) found that herbivory removal enhanced the population growth of the perennial herb *Panax quinquefolius* in an old-growth beech and maple forest; Bermingham (2010) demonstrated that exclusion of browsing accelerated the population growth of rare plant *Polemonium vanbruntiae* in minerotrophic wetlands of the Green Mountain National Forest. Similar positive effects of grazing removal have also been observed for shrubs and trees (Hunt 2001; Aschero et al. 2016). These findings, together with our results, suggest that the grazing removal may be a consistently effective grazing management action for biodiversity conservation.

The impacts of grazing removal on population viability are largely depended on climate warming. In our study, warming reduced the population growth rate of both study species under grazing removal. Similarly, Evju et al. (2010) reported that elevated temperatures accelerated the population decline of the small herb *Viola biflora* following cessation of grazing in an alpine habitat. Williams et al. (2007) found that four years warming by 2□ reduced the population growth of four perennial plant co-existing species under fencing conditions in southeastern Australian. These findings, together with our results, suggest that climate warming is generally detrimental to population maintenance under grazing removal. Importantly, we found that the negative effects of climate warming on population growth under grazing removal outweighed the positive effects of grazing removal alone. This suggests grazing removal may be insufficient to rescue plant species in danger if warming trends continue. Taken together, these findings suggest that successful management strategies must fully consider the impact of climate warming on population dynamics of plant species.

The impact of grazing removal on population viability under ambient and warmed conditions may be modulated by altitude, altering both the magnitude and direction of demographic responses. In our results, we found that the minor negative effect of grazing removal on the population growth rate of *C. atrofusca* was reversed at higher altitude, whereas the positive effect of grazing removal on the population growth rate of *S. sibiricum* weakened at higher altitude. Such contrasting altitude-dependent demographic responses under grazing removal have been reported in several studies (Jónsdóttir 1991; Miller et al. 2009). For example, Jónsdóttir (1991) reported that negative effect of grazing removal on the population growth rate of the clonal sedge *Carex bigelowii* was alleviated at higher altitude, while Miller *et al*. (2009) reported the positive effect of excluding insect herbivores on the population growth of the perennial plant *Opuntia imbricat* diminished at higher altitude. On the other hand, our results showed that the negative effect of warming on the population growth rate under grazing removal was alleviated at higher altitude for both species. This pattern may reflect population-specific adaptations to local climate (Peterson et al. 2018), leading to weaker population responses to warming at higher altitude. Collectively, these findings highlight the critical role of altitude in shaping species’ demographic responses to grazing and warming.

Grazing removal may impact population viability by differentially affecting demographic processes. Demographic compensation among vital rates may allow species to maintain viable populations under grazing removal (Villellas et al. 2015; Sheth & Angert 2018). In our study, grazing removal-induced reduction in survival has a substantial negative contribution to population growth in both species under both ambient and warmed conditions. Compensatory increases in contributions from growth and shrinkage offset these reductions, thereby benefiting the population growth in both species. Demographic compensation under grazing removal has also been documented in previous studies (Farrington et al. 2009; Evju et al. 2010, 2011; Johansen et al. 2016). For example, Evju et al. (2010) reported the reduced growth under grazing removal was largely compensated by increases in fecundity and stasis, thereby alleviating the population decline under grazing removal; Evju et al. (2011) reported that decreased contributions from retrogression and clonal reproduction were completely compensated by increased contributions from survival, thereby stabilizing population growth of the clonal herb *Geranium sylvaticum* under grazing removal; Farrington et al. (2019) found the decreased contributions in regression were overcompensated by increases in fecundity and stasis, thereby enhancing the population growth of the perennial herb *Panax quinquefolius* under grazing removal. Nevertheless, climate warming caused a breakdown of such demographic compensation in populations under grazing removal, leading to population declines in both study species. This finding underscores that demographic resilience to grazing removal can fail under warming, highlighting the need to consider interactive effects of grazing activities and climate change in conservation strategies.

### Adaptive grazing management implications under contemporary conditions and future warming

Livestock grazing is a dominant land use practice in alpine grassland ecosystems (Wang et al. 2022; Zhang et al. 2023). Nevertheless, decades of extensive overgrazing accelerated the grassland degradation on the Tibetan Plateau (Bardgett et al. 2021; Wang et al. 2022). Grazing removal has been widely implemented in the region as a conservation and restoration strategy (Sun et al. 2020; Wang et al. 2022). Our study showed that current grazing removal can maintain or enhance population maintenance of both study species, indicating that short-term grazing removal (< 10 years) can help stabilize plant populations in degraded grassland on the Tibetan Plateau. However, the positive effects of grazing removal on population growth were nullified under warming in both study species, implying that current grazing management is unsustainable under future warming scenario. Therefore, adjusting grazing regime could help rescue warming-induced population declines (Andrello et al. 2012; Miao et al. 2025). Further research is needed to determine the optimal grazing intensity and season to maintain viable population under climate change.

Effective grazing management strategies should target survival, growth, and shrinkage in plant species. In our study, survival was not only key for population maintenance of both species, but also the main demographic rate limiting population growth under grazing removal and warming, suggesting that grazing activities should be regulated to minimize mortality. In addition, increased growth and reduced shrinkage under grazing removal made considerable positive contributions to the population growth of both species in the current study. Previous studies on the Tibetan Plateau have reported that grazing removal significantly increased the average plant size in terms of height and aboveground biomass (Yang et al. 2016; Ji et al. 2020; Zhang et al. 2023). Therefore, these findings together with our results suggest that grazing activities should also be managed to prevent substantial reductions in growth and increases in shrinkage. Given survival, growth and shrinkage are relatively easy to monitor, they could potentially serve as warning signals for the management of alpine grasslands under climate change.

Communities at high latitudes and elevations, like the Tibetan plateau system studied here, serve as bellwethers for global change, especially in response to grazing activities and climate warming. We show that current grazing removal generally stabilizes or enhances the population growth of alpine species, whereas long-term warming leads to marked population declines. This highlights the urgent need to adjust grazing regimes in the context of climate change. Demographic compensation among vital rates can promote population maintenance under grazing removal; however, warming can disrupt this mechanism. Monitoring key demographic rates— particularly survival, growth, and shrinkage—particularly survival, growth and shrinkage, could provide predictive metrics to guide adaptive management. Overall, these findings emphasize that carefully calibrated grazing regimes, informed by demographic monitoring, may help maintain the long-term viability of alpine plant communities under future climate scenarios.

## Acknowledgements

This work was funded by the National Natural Science Foundation of China (31971423) and the cost-share exchange projects between the National Natural Science Foundation of China and the Royal Society of the United Kingdom (32011530169).

## Author contributions

SLL and JSH conceived the study. HTM and XW collected the data, HTM constructed models and wrote first draft of the manuscript, and all authors contributed substantially to revisions. All authors approved the final manuscript.

## Data availability

Data and code are archived in Zenodo (https://doi.org/10.5281/zenodo.17623384) and are available for peer review via the following private link: https://zenodo.org/records/17623384?token=eyJhbGciOiJIUzUxMiJ9.eyJpZCI6Ijc0ZmE5NDA2LWNhODEtNDA5ZC1iZTg2LTFkYjQ2NjRlOWFjZSIsImRhdGEiOnt9LCJyYW5kb20iOiIzNjA0MzRlYmIwNzllMTMxZDMwMDVhZWFlMGMzYzM1NCJ9.ri7962HFl8znsGf8qCeojZ4cRgPuUfkUVRYVOiJpEEkEXl10mupwGCt_79CfCXdrGrVHy4E2NY_Sz3U3i79dow.

## References

1. Andrello, M., Bizoux, J.P., Barbet-Massin, M., Gaudeul, M., Nicole, F. & Till-Bottraud, I. (2012). Effects of management regimes and extreme climatic events on plant population viability in Eryngium alpinum. Biol. Conserv., 147, 99–106.

2. Andrello, M., de Villemereuil, P., Carboni, M., Busson, D., Fortin, M.J., Gaggiotti, O.E. et al. (2020). Accounting for stochasticity in demographic compensation along the elevational range of an alpine plant. Ecol. Lett., 23, 870–880.

3. Aschero, V., Morris, W.F., Vazquez, D.P., Alvarez, J.A. & Villagra, P.E. (2016). Demography and population growth rate of the tree Prosopis flexuosa with contrasting grazing regimes in the Central Monte Desert. For. Ecol. Manage., 369, 184–190.

4. Bardgett, R.D., Bullock, J.M., Lavorel, S., Manning, P., Schaffner, U., Ostle, N. et al. (2021). Combatting global grassland degradation. Nat. Rev. Earth Environ., 2, 720–735.

5. Bates, D., Mächler, M., Bolker, B. & Walker, S. (2015). Fitting linear mixed-efects models using lme4. J. Stat. Softw., 67, 1–48.

6. Bermingham, L.H. (2010). Deer herbivory and habitat type influence long-term population dynamics of a rare wetland plant. Plant Ecol., 210, 359–378.

7. Brooks, M.E., Kristensen, K., van Benthem, K.J., Magnusson, A., Berg, C.W., Nielsen, A. et al. (2017). glmmTMB balances speed and flexibility among packages for zero-inflated generalized linear mixed modeling. R J., 9, 378–400.

8. Burnham, K.P. & Anderson, D.R. (2004). Multimodel inference - understanding AIC and BIC in model selection. Sociol. Methods Res., 33, 261–304.

9. Cribari-Neto, F. & Zeileis, A. (2010). Beta regression in R. J. Stat. Softw., 34, 1–24.

10. Davison, R., Nicole, F., Jacquemyn, H. & Tuljapurkar, S. (2013). Contributions of covariance: decomposing the components of stochastic population growth in Cypripedium calceolus. Am. Nat., 181, 410–420.

11. Davison, R., Stadman, M. & Jongejans, E. (2019). Stochastic effects contribute to population fitness differences. Ecol. Modell., 408, 108760.

12. de Kroon, H., van Groenendael, J. & Ehrlen, J. (2000). Elasticities: a review of methods and model limitations. Ecology, 81, 607–618.

13. Easterling, M.R., Ellner, S.P. & Dixon, P.M. (2000). Size-specific sensitivity: Applying a new structured population model. Ecology, 81, 694–708.

14. Elderd, B.D. & Doak, D.F. (2006). Comparing the direct and community-mediated effects of disturbance on plant population dynamics: flooding, herbivory and Mimulus guttatus. J. Ecol., 94, 656–669.

15. Elderd, B.D. & Miller, T.E.X. (2016). Quantifying demographic uncertainty: bayesian methods for integral projection models. Ecol. Monogr., 86, 125–144.

16. Ellner, S.P., Childs, D.Z. & Rees, M. (2016). Data-driven modelling of structured populations: a practical guide to the integral projection model. Springer International Publishing, Switzerland.

17. Ellner, S.P. & Rees, M. (2006). Integral projection models for species with complex demography. American Naturalist, 167, 410–428.

18. Elmendorf, S.C., Henry, G.H.R., Hollister, R.D., Björk, R.G., Boulanger-Lapointe, N., Cooper, E.J. et al. (2012). Plot-scale evidence of tundra vegetation change and links to recent summer warming. Nat. Clim. Change, 2, 453–457.

19. Elsen, P.R. & Tingley, M.W. (2015). Global mountain topography and the fate of montane species under climate change. Nat. Clim. Change, 5, 772–U192.

20. Evers, S.M., Knight, T.M., Inouye, D.W., Miller, T.E.X., Salguero-Gómez, R., Iler, A.M. et al. (2021). Lagged and dormant season climate better predict plant vital rates than climate during the growing season. Glob. Change Biol., 27, 1927–1941.

21. Evju, M., Halvorsen, R., Rydgren, K., Austrheim, G. & Mysterud, A. (2010). Interactions between local climate and grazing determine the population dynamics of the small herb Viola biflora. Oecologia, 163, 921–933.

22. Evju, M., Halvorsen, R., Rydgren, K., Austrheim, G. & Mysterud, A. (2011). Effects of sheep grazing and temporal variability on population dynamics of the clonal herb Geranium sylvaticum in an alpine habitat. Plant Ecol., 212, 1299–1312.

23. Farrington, S.J., Muzika, R.M., Drees, D. & Knight, T.M. (2009). Interactive effects of harvest and deer herbivory on the population dynamics of American ginseng. Conserv. Biol., 23, 719–728.

24. Fox, J. & Weisberg, S. (2019). An R companion to applied regression. Third edn. SAGE Publications, Inc, Christopher Hare.

25. Gao, J.J. & Carmel, Y. (2020). A global meta-analysis of grazing effects on plant richness. Agr. Ecosyst. Environ., 302, 107072.

26. Golodets, C., Kigel, J. & Sternberg, M. (2010). Recovery of plant species composition and ecosystem function after cessation of grazing in a Mediterranean grassland. Plant Soil, 329, 365–378.

27. Hejcman, M., Hejcmanová, P., Pavlu, V. & Benes, J. (2013). Origin and history of grasslands in Central Europe - a review. Grass Forage Sci., 68, 345–363.

28. Hunt, L.P. (2001). Heterogeneous grazing causes local extinction of edible perennial shrubs: a matrix analysis. J. Appl. Ecol., 38, 238–252.

29. IPCC (2023). Sections. In: Climate Change 2023: Synthesis Report. Contribution of Working Groups I, II and III to the Sixth Assessment Report of the Intergovernmental Panel on Climate Change, IPCC, Geneva, Switzerland.

30. Ji, L., Qin, Y., Jimoh, S.O., Hou, X.Y., Zhang, N., Gan, Y.M. et al. (2020). Impacts of livestock grazing on vegetation characteristics and soil chemical properties of alpine meadows in the eastern Qinghai-Tibetan Plateau. Ecoscience, 27, 107–118.

31. Johansen, L., Wehn, S. & Hovstad, K.A. (2016). Clonal growth buffers the effect of grazing management on the population growth rate of a perennial grassland herb. Flora, 223, 11–18.

32. Jónsdóttir, I.S. (1991). Effects of grazing on tiller size and population dynamcis in a clonal sedge (Carex bigelowii). Oikos, 62, 177–188.

33. Knight, T.M. (2004). The effect of herbivory and pollen limitation on a declining population of Trillium grandiflorum. Ecol. Appl., 14, 915–928.

34. Körner, C. (2007). The use of ‘altitude’ in ecological research. Trends Ecol. Evol., 22, 569–574.

35. Larios, L. & Hallett, L.M. (2022). Incorporating temporal dynamics to enhance grazing management outcomes for a long-lived species. J. Appl. Ecol., 59, 2936–2946.

36. Li, S.L., Yu, F.H., Werger, M.J.A., Dong, M., During, H.J. & Zuidema, P.A. (2015). Mobile dune fixation by a fast-growing clonal plant: a full life-cycle analysis. Sci. Rep., 5, 8935.

37. Li, S.L., Yu, F.H., Werger, M.J.A., Dong, M., Ramula, S. & Zuidema, P.A. (2013). Understanding the effects of a new grazing policy: the impact of seasonal grazing on shrub demography in the Inner Mongolian steppe. J. Appl. Ecol., 50, 1377–1386.

38. Maron, J.L., Baer, K.C. & Angert, A.L. (2014). Disentangling the drivers of context-dependent plant-animal interactions. J. Ecol., 102, 1485–1496.

39. Martorell, C. (2007). Detecting and managing an overgrazing-drought synergism in the threatened Echeveria longissima (Crassulaceae): the role of retrospective demographic analysis. Popul. Ecol., 49, 115–125.

40. Menges, E.S. (2000). Population viability analyses in plants: challenges and opportunities. Trends Ecol. Evol., 15, 51–56.

41. Miao, H.T., Salguero-Gómez, R., Shea, K., Keller, J.A., Zhang, Z.H., He, J.S. et al. (2025). Differences in adult survival drive divergent demographic responses to warming on the Tibetan Plateau. Ecology, 106, e4533.

42. Miller, T.E.X., Louda, S.M., Rose, K.A. & Eckberg, J.O. (2009). Impacts of insect herbivory on cactus population dynamics: experimental demography across an environmental gradient. Ecol. Monogr., 79, 155–172.

43. Nomoto, H.A. & Alexander, J.M. (2021). Drivers of local extinction risk in alpine plants under warming climate. Ecol. Lett., 24, 1157–1166.

44. Peterson, M.L., Doak, D.F. & Morris, W.F. (2018). Both life-history plasticity and local adaptation will shape range-wide responses to climate warming in the tundra plant Silene acaulis. Glob. Change Biol., 24, 1614–1625.

45. Prager, C.M., Classen, A.T., Sundqvist, M.K., Barrios-Garcia, M.N., Cameron, E.K., Chen, L.T. et al. (2022). Integrating natural gradients, experiments, and statistical modeling in a distributed network experiment: an example from the WaRM Network. Ecol. Evol., 12, e9396.

46. Rees, M. & Ellner, S.P. (2009). Integral projection models for populations in temporally varying environments. Ecol. Monogr., 79, 575–594.

47. Schonswetter, P., Popp, M. & Brochmann, C. (2006). Central Asian origin of and strong genetic differentiation among populations of the rare and disjunct Carex atrofusca (Cyperaceae) in the Alps. J. Biogeogr., 33, 948–956.

48. Schrama, M., Quist, C.W., de Groot, G.A., Cieraad, E., Ashworth, D., Laros, I. et al. (2023). Cessation of grazing causes biodiversity loss and homogenization of soil food webs. Proc. R. Soc. B, 290, 20231345.

49. Seddon, A.W.R., Macias-Fauria, M., Long, P.R., Benz, D. & Willis, K.J. (2016). Sensitivity of global terrestrial ecosystems to climate variability. Nature, 531, 229–+.

50. Sheth, S.N. & Angert, A.L. (2018). Demographic compensation does not rescue populations at a trailing range edge. Proc. Natl. Acad. Sci. USA, 115, 2413–2418.

51. Sletvold, N., Dahlgren, J.P., Oien, D.-I., Moen, A. & Ehrlen, J. (2013). Climate warming alters effects of management on population viability of threatened species: results from a 30-year experimental study on a rare orchid. Glob. Change Biol., 19, 2729–2738.

52. Struckman, S., Couture, J.J., LaMar, M.D. & Dalgleish, H.J. (2019). The demographic effects of functional traits: an integral projection model approach reveals population-level consequences of reproduction-defence trade-offs. Ecol. Lett., 22, 1396–1406.

53. Sun, J., Liu, M., Fu, B.J., Kemp, D., Zhao, W.W., Liu, G.H. et al. (2020). Reconsidering the efficiency of grazing exclusion using fences on the Tibetan Plateau. Sci. Bull., 65, 1405–1414.

54. Sun, Y.X., Liu, S.L., Liu, Y.X., Dong, Y.H., Li, M.Q., An, Y. et al. (2022). Grazing intensity and human activity intensity data sets on the Qinghai-Tibetan Plateau during 1990-2015. Geosci. Data J., 9, 140–153.

55. Tenhumberg, B., Crone, E.E., Ramula, S. & Tyre, A.J. (2018). Time-lagged effects of weather on plant demography: drought and Astragalus scaphoides. Ecology, 99, 915–925.

56. Tredennick, A.T., Teller, B.J., Adler, P.B., Hooker, G. & Ellner, S.P. (2018). Size-by-environment interactions: a neglected dimension of species’ responses to environmental variation. Ecol. Lett., 21, 1757–1770.

57. Tuljapurkar, S. (1990). Population Dynamics in Variable Environments. Springer, New York.

58. van de Pol, M., Bailey, L.D., McLean, N., Rijsdijk, L., Lawson, C.R. & Brouwer, L. (2016). Identifying the best climatic predictors in ecology and evolution. Methods Ecol. Evol., 7, 1246–1257.

59. Verburg, R.W., Kwant, R. & Werger, M.J.A. (1996). The effect of plant size on vegetative reproduction in a pseudo-annual. Vegetatio, 125, 185–192.

60. Verdoodt, A., Mureithi, S.M., Ye, L.M. & Van Ranst, E. (2009). Chronosequence analysis of two enclosure management strategies in degraded rangeland of semi-arid Kenya. Agr. Ecosyst. Environ., 129, 332–339.

61. Villellas, J., Doak, D.F., Garcia, M.B. & Morris, W.F. (2015). Demographic compensation among populations: what is it, how does it arise and what are its implications? Ecol. Lett., 18, 1139–1152.

62. Wang, H., Liu, H., Cao, G., Ma, Z., Li, Y., Zhang, F. et al. (2020). Alpine grassland plants grow earlier and faster but biomass remains unchanged over 35 years of climate change. Ecol. Lett., 23, 701–710.

63. Wang, Y., Lv, W., Xue, K., Wang, S., Zhang, L., Hu, R. et al. (2022). Grassland changes and adaptive management on the Qinghai-Tibetan Plateau. Nat. Rev. Earth Environ., 3, 668–683.

64. Watkinson, A.R. & Ormerod, S.J. (2001). Grasslands, grazing and biodiversity: editors’ introduction. J. Appl. Ecol., 38, 233–237.

65. Williams, A.L., Wills, K.E., Janes, J.K., Schoor, J.K.V., Newton, P.C.D. & Hovenden, M.J. (2007). Warming and free-air CO2 enrichment alter demographics in four co-occurring grassland species. New Phytol., 176, 365–374.

66. Xiang, M.X., Wu, J.X., Duo, L., Niu, B. & Zhang, X.Z. (2023). The effects of grazing and fencing on grassland productivity and diversity in alpine grassland ecosystem in the Tibetan highland. Global Ecol. Conserv., 44, e02495.

67. Xu, L.H., Cao, G.X., Wang, Y.N., Hao, J., Wang, Y.H., Yu, P.T. et al. (2020). Components of stand water balance of a larch plantation after thinning during the extremely wet and dry years in the Loess Plateau, China. Global Ecol. Conserv., 24, e01307.

68. Yang, Z.a., Xiong, W., Xu, Y., Jiang, L., Zhu, E., Zhan, W. et al. (2016). Soil properties and species composition under different grazing intensity in an alpine meadow on the eastern Tibetan Plateau, China. Environ. Monit. Assess., 188, 678.

69. Yang, Z.H. (1998). Likelihood ratio tests for detecting positive selection and application to primate lysozyme evolution. Mol. Biol. Evol., 15, 568–573.

70. Yao, T.D., Bolch, T., Chen, D.L., Gao, J., Immerzeel, W., Piao, S. et al. (2022). The imbalance of the Asian water tower. Nat. Rev. Earth Environ., 3, 618–632.

71. Zhang, Z., Zhao, Y., Lin, H., Li, Y., Fu, J., Wang, Y. et al. (2023). Comprehensive analysis of grazing intensity impacts alpine grasslands across the Qinghai-Tibetan Plateau: A meta-analysis. Front. Plant Sci., 13, 1083709.

72. Zhong, L., Ma, Y., Xue, Y. & Piao, S. (2019). Climate Change Trends and Impacts on Vegetation Greening Over the Tibetan Plateau. J. Geophys. Res.: Atmos., 124, 7540–7552.

